# ATP synthesis of *Enterococcus hirae* V-ATPase driven by sodium motive force

**DOI:** 10.1101/2024.12.26.630461

**Authors:** Akihiro Otomo, Lucy Gao Hui Zhu, Yasuko Okuni, Mayuko Yamamoto, Ryota Iino

**Author notes:** Corresponding authors :Akihiro Otomo, Ryota Iino. Present address: Department of Chemistry, Graduate School of Science, Kyoto University, Kitashirakawa-Oiwakecho, Sakyo-ku, Kyoto 606–8502, Japan (Akihiro Otomo).

## Abstract

V-ATPases generally function as ion pumps driven by ATP hydrolysis in the cell, but their capability of ATP synthesis remains largely unexplored. Here we show ATP synthesis of Na^+^-transporting *Enterococcus hirae* V-ATPase (EhV_o_V_1_) driven by electrochemical potential gradient of Na^+^ across the membrane (sodium motive force, *smf*). We reconstituted EhV_o_V_1_ into liposome and performed a luciferin/luciferase-based assay to analyze ATP synthesis quantitatively. Our result demonstrates that EhV_o_V_1_ synthesizes ATP with a rate of 4.7 s^-1^ under high *smf* (269.3 mV). The Michaelis constants for ADP (21 µM) and inorganic phosphate (2.1 mM) in ATP synthesis reaction were comparable to those for ATP synthases, suggesting similar substrate affinities among rotary ATPases regardless of their physiological functions. Both components of *smf*, Na^+^ concentration gradient across the membrane (ΔpNa) and membrane potential (Δψ), contributed to ATP synthesis, with ΔpNa showing a slightly larger impact. At the equilibrium points where *smf* and Gibbs free energy of ATP synthesis are balanced, EhV_o_V_1_ showed reversible reactions between ATP synthesis and hydrolysis. The obtained Na^+^/ATP ratio (3.2 ± 0.4) closely matched the value expected from the structural symmetry ratio between EhV_o_ and EhV_1_ (10/3 = 3.3), indicating tight coupling between ATP synthesis/hydrolysis and Na^+^ transport. These results reveal inherent functional reversibility of EhV_o_V_1_. We propose that physiological function of EhV_o_V_1_ *in vivo* is determined by relatively small *smf* against large Gibbs free energy of ATP synthesis, in addition to the absence of inhibitory mechanisms of ATP hydrolysis which are known for ATP synthases.

## Introduction

F-, V-, and A-ATPases (F_o_F_1_, V_o_V_1_, and A_o_A_1_, respectively) are rotary motor proteins and classified together as rotary ATPases (1–4). These rotary ATPases share a common structural architecture: a cytoplasmic domain (F_1_, V_1_ or A_1_) for ATP synthesis and hydrolysis and a membrane-embedded domain (F_o_, V_o_, or A_o_) for transport of protons (H^+^) or sodium ion (Na^+^). F-ATPases are present in mitochondria and chloroplasts of eukaryotes and plasma membranes of many bacteria, and primarily function as ATP synthases. ATP synthesis reaction is driven by the ion motive force (*imf*), the electrochemical potential gradient of the ion across the membrane, generated by respiratory chains or photosynthesis. F-ATPases show reversible energy conversion, being capable of pumping H^+^ or Na^+^ by hydrolyzing ATP when the *imf* is insufficient and/or the ATP/ADP ratio is high in the cell (1).

V-ATPases are located in eukaryotic organelles and some bacterial plasma membranes, and generally operate as ATP hydrolysis-driven ion pumps. Bacterial V-ATPases consist of A, B, D, E, F, G, d subunits in V_1_ and a, c subunits in V_o_, and eukaryotic V-ATPases often have additional, organism-specific subunits and isoforms (5). V-ATPases play crucial roles in organelle acidification, homeostasis of H^+^ or Na^+^ concentration in the cell, and other cellular processes (6, 7). Interestingly, the V/A-ATPase from *Thermus thermophilus* (TtV_o_V_1_) functions as an ATP synthase in the cell, in contrast to other V-ATPases (8, 9). Archaea possess A-ATPases structurally similar to V-ATPases but primarily function as ATP synthases, highlighting the diverse roles of these rotary ATPases across life domains (3, 4).

ATP synthesis catalyzed by rotary ATPases has been mostly studied with F- and A- ATPases and TtV_o_V_1_, all of which physiologically function as ATP synthases (10–17). Very few studies have been conducted with V-ATPases, so our understanding of their potential reversibility is very limited. Hirata et al., provided first evidence for ATP synthesis of yeast V-ATPase in vacuolar membrane driven by the *imf* generated by a pyrophosphatase, and successfully estimated the Michaelis constant (*K*_m_) for ADP and the maximum velocity (18). However, for V-ATPases, important parameters for reversibility remain unexplored, such as equilibrium points between ATP synthesis and hydrolysis, and ion/ATP coupling ratio between V_o_ and V_1_.

The ion/ATP coupling ratio is one of the most crucial parameters for rotary ATPases, which determines the equilibrium points and directions of their operations, ATP synthesis or hydrolysis (11). This ratio represents the number of ions transported per ATP molecule synthesized or hydrolyzed. The rotary catalysis in F_1_, V_1_, and A_1_ accompanies the rotation of the central stalk in the hexameric stator ring. Single rotation is coupled with synthesis or hydrolysis of 3 ATP molecules, reflecting the three catalytic sites in the hexameric stator ring. The rotation of the central stalk in F_1_, V_1_, and A_1_ is coupled with ion transport through a stator a-subunit and a rotor c-ring of F_o_, V_o_, and A_o_ (1, 2). The c-ring, composed of multiple c-subunits each with one ion binding site, determines the number of ions transported per single rotation. The number of c-subunits in a c-ring varies among species (8 to 17), resulting in ion/ATP coupling ratio ranging from 2.7 to 5.7 (19). The direction of the chemical reaction, ATP synthesis or hydrolysis in these enzymes is thermodynamically governed by the relationship between the Gibbs free energy of ATP synthesis and *imf*:

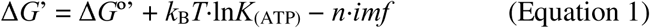

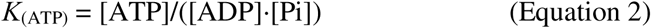

where Δ*G*°’ is the standard Gibbs free energy of ATP synthesis, *k*_B_ is the Boltzmann constant, *T* is absolute temperature, *K*_(ATP)_ is the ratio of ATP concentration ([ATP]) to ADP and inorganic phosphate (Pi) concentrations ([ADP] and [Pi], respectively), and *n* is the ion/ATP coupling ratio, respectively. A large *n* value allows ATP synthesis even at low *imf*, and this value is presumably related to the environments in which living organisms grow (12, 13, 19, 20).

The V-ATPase from *Enterococcus hirae* (EhV_o_V_1_) is an ion pump that transports Na^+^, enabling *E. hirae* to grow in alkaline environments by maintaining Na^+^ homeostasis (21, 22). Extensive studies have elucidated the structure of EhV_o_V_1_ (23–28) and its rotary dynamics driven by ATP hydrolysis (29–32). The EhV_o_ has a c-ring composed of 10 c-subunits (28). Therefore, expected Na^+^/ATP ratio for EhV_o_V_1_ is 3.3 (10/3), similar to H^+^/ATP ratio of 3.3 for ATP synthases from *Escherichia coli* (33) and thermophilic *Bacillus* PS3 (34–36). However, no studies have examined whether EhV_o_V_1_ can synthesize ATP using the sodium motive force (*smf*), composed of the electrochemical potential of Na^+^ across the membrane (ΔpNa) and the membrane potential (Δψ).

The present study aims to elucidate the ATP synthesis capability of EhV_o_V_1_ driven by the *smf*. We reconstituted EhV_o_V_1_ into liposomes, allowing precise control of the *smf*, and employed a luciferin/luciferase-based assay for ATP detection (Figure 1). This approach enabled us to demonstrate that EhV_o_V_1_ can synthesize ATP driven by *smf*. Quantitative analysis revealed [ADP], [Pi], and *smf* dependences of ATP synthesis rate, and kinetic contributions of ΔpNa and Δψ to ATP synthesis. Furthermore, we revealed high thermodynamic efficiency (3.2 Na^+^/ATP ratio comparable to the value of 3.3 expected from the structure) of EhV_o_V_1_ at the equilibrium point between ATP synthesis and hydrolysis, similar to previously studied ATP synthases (10, 37). Our results provide new insights into the inherent functional reversibility of V-ATPases, and raise intriguing questions about the physiological relevance of their ATP synthesis capability.

**Figure 1.**
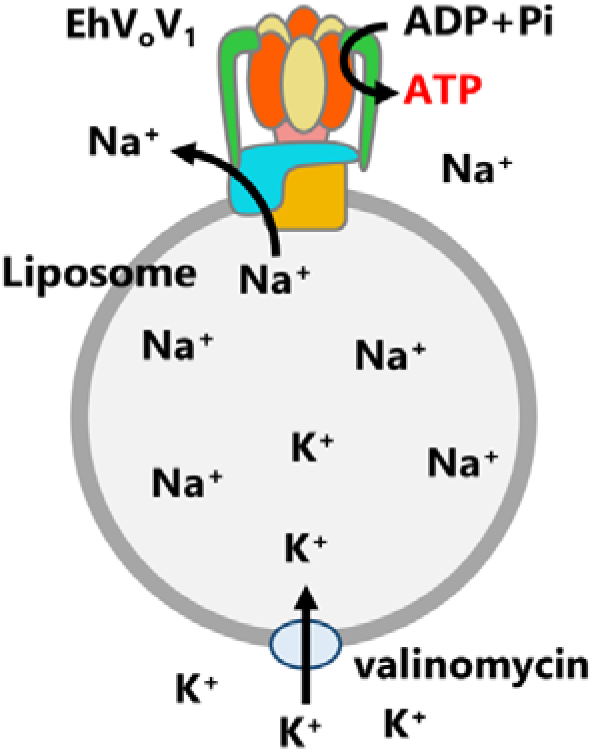
Schematic illustration of ATP synthesis experiment of EhV_o_V_1_ reconstituted into liposome. ATP synthesis was driven by *smf*, generated by sum of Na^+^ concentration gradient (ΔpNa) and K^+^- valinomycin diffusion potential (Δψ) across the membrane. Synthesized ATP was detected as luminescence using luciferin/luciferase system.

## Results

### Demonstration of smf-driven ATP synthesis of EhV_o_V_1_ and its dependence on [ADP] and [Pi]

The ATP synthesis activity of EhV_o_V_1_ reconstituted into liposome was measured via the luminescence intensity change of luciferin/luciferase system at 25°C (Figure 1). The imposed *smf* was calculated by the equation:

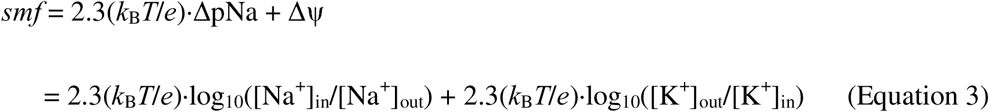

where *e* is the elementary charge, and [Na^+^]_in_ and [Na^+^]_out_ ([K^+^]_in_ and [K^+^]_out_) are Na^+^ (K^+^) concentrations inside and outside the liposome, respectively. The Δψ was applied using K^+^- valinomycin diffusion potential. To accurately calculate *smf*, we measured ion concentrations unintendedly contained in the buffers and regents using inductively coupled plasma optical emission spectrometer (ICP-OES) (Figure S1). The solution compositions used for measurements in this study are summarized in Table S1-S3.

Figure 2A shows the time courses of ATP synthesis of EhV_o_V_1_ under the high *smf* (269.3 mV: Δψ of 154.7 mV and 2.3(*k*_B_*T*/*e*)·ΔpNa of 114.6 mV) and various [ADP] and [Pi]. Note that to facilitate ATP synthesis, we did not add ATP intentionally but a small amount of ATP was contaminated in ADP (<0.003%) (Table S1). The observed increments in the luminescence intensity after the addition of PL indicate that EhV_o_V_1_ synthesizes ATP driven by *smf*. The intensity changes were converted into [ATP] changes based on the luminescence increase caused by the addition of 200 nM ATP at the end of each measurement. ATP synthesis rates were estimated by fitting the intensity change after the addition of PL with a single exponential function and calculating the initial slope of the fitted curve.

**Figure 2.**
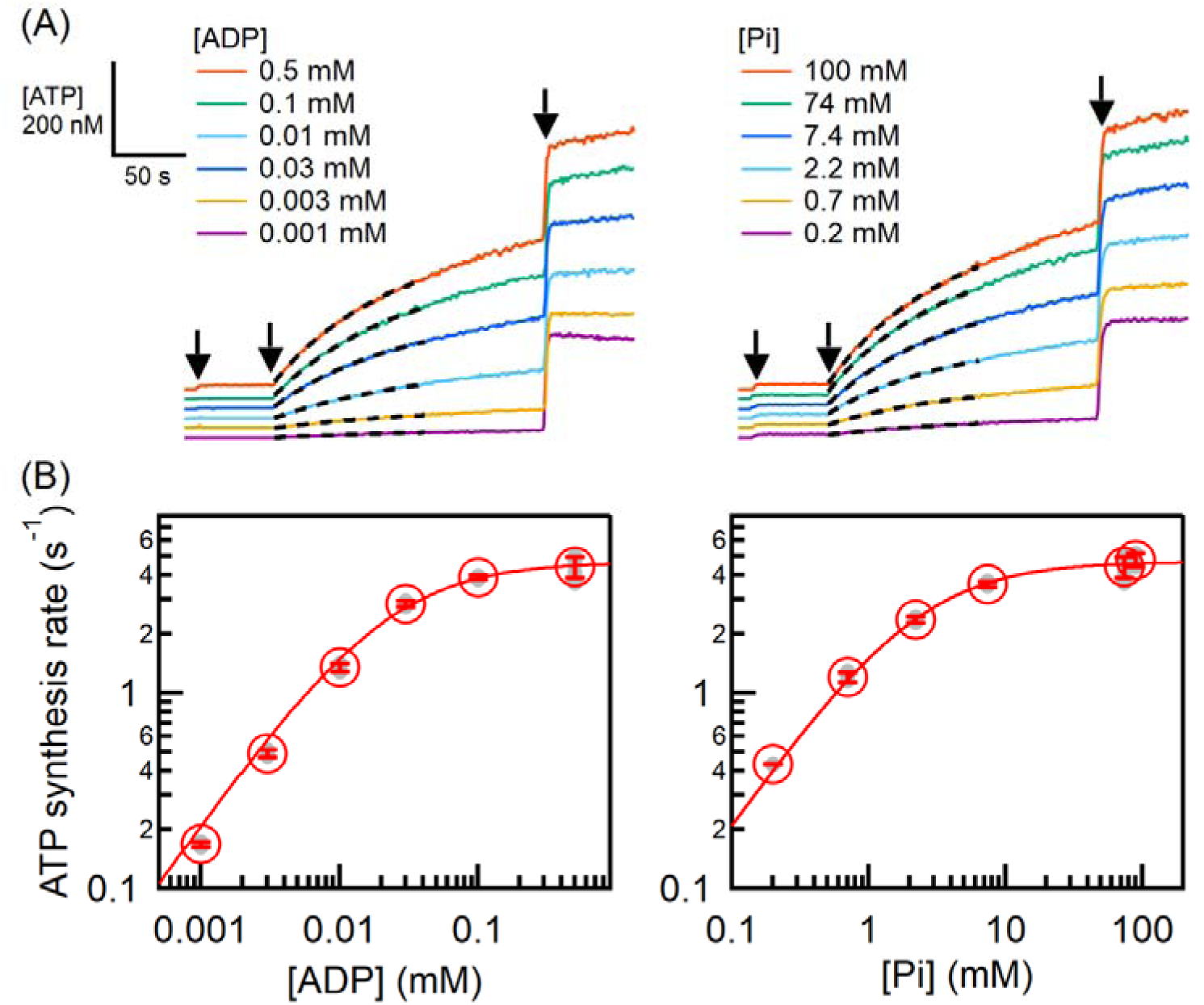
ATP synthesis of EhV_o_V_1_ driven by *smf*. (A) [ADP] (left) and [Pi] (right) dependences of ATP synthesis activity of EhV_o_V_1_ under *smf* of 269.3 mV (Δψ of 154.7 mV and 2.3(*k*_B_*T*/*e*)·ΔpNa of 114.6 mV). [Pi] (left) and [ADP] (right) were set at 74 and 0.5 mM, respectively. ATP was not added but contaminated in ADP (<0.003%) (Table S1). ADP (0.001 - 0.5 mM), PL, and ATP (200 nM) were added at 10, 60, and 240 sec, respectively, as indicated by black arrows. The black dashed lines represent the fitting with a single exponential function for Δt = 100 sec after the addition of PL. (B) ATP synthesis rate against [ADP] (left) and [Pi] (right). Red open circles represent mean values obtained from more than three measurements. Individual data points are shown as gray filled circles. Error bars indicate standard deviations. Red lines show the fitting with the Michaelis-Menten equation.

As a result, [ADP] and [Pi] dependences of the ATP synthesis rates were well fitted by the Michaelis-Menten equation, yielding the same *k*_cat_ of 4.7 s^-1^ for both [ADP] and [Pi] dependences, and *K*_m, ADP_ = 21 µM and *K*_m, Pi_ = 2.1 mM for [ADP] and [Pi] dependences, respectively (Figure 2B). These *K*_m_ values for EhV_o_V_1_ were comparable to those of other rotary ATPases that function as ATP synthases physiologically (Table 1). The comparable values of *K*_m_ suggest that the affinities of ADP and Pi in ATP synthesis reaction are similar across EhV_o_V_1_ and these enzymes regardless of their primary physiological functions.

**Table 1.**
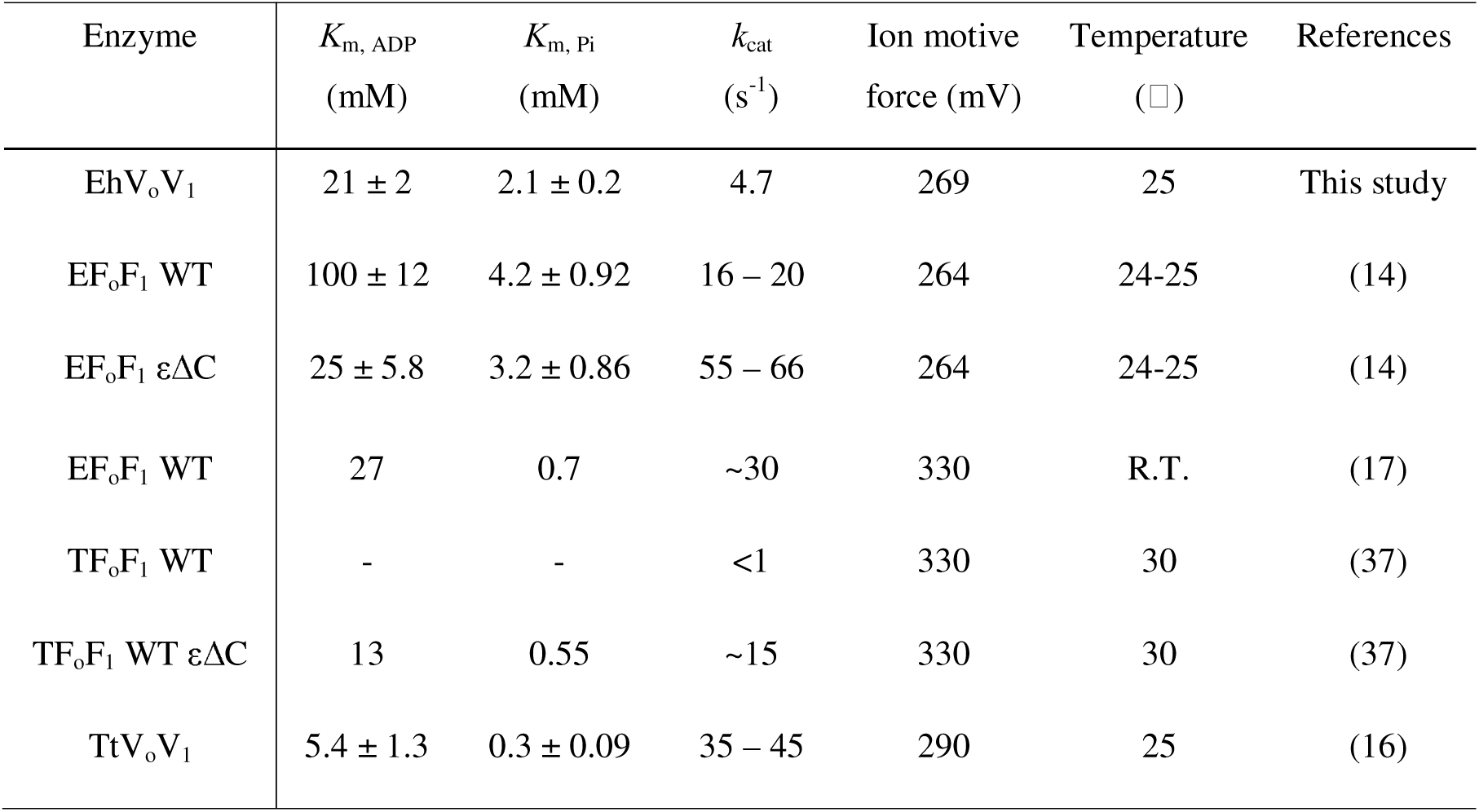
Kinetic parameters for ATP synthesis of EhV_o_V_1_ and other rotary ATPases.

| Enzyme | $K_{m, \text{ADP}}$<br>(mM) | $K_{m, \text{Pi}}$<br>(mM) | $k_{\text{cat}}$<br>(s<sup>-1</sup>) | Ion motive<br>force (mV) | Temperature<br>(°C) | References |
| --- | --- | --- | --- | --- | --- | --- |
| EhV<sub>o</sub>V<sub>1</sub> | 21 ± 2 | 2.1 ± 0.2 | 4.7 | 269 | 25 | This study |
| EF<sub>o</sub>F<sub>1</sub> WT | 100 ± 12 | 4.2 ± 0.92 | 16 – 20 | 264 | 24-25 | (14) |
| EF<sub>o</sub>F<sub>1</sub> εΔC | 25 ± 5.8 | 3.2 ± 0.86 | 55 – 66 | 264 | 24-25 | (14) |
| EF<sub>o</sub>F<sub>1</sub> WT | 27 | 0.7 | ~30 | 330 | R.T. | (17) |
| TF<sub>o</sub>F<sub>1</sub> WT | - | - | <1 | 330 | 30 | (37) |
| TF<sub>o</sub>F<sub>1</sub> WT εΔC | 13 | 0.55 | ~15 | 330 | 30 | (37) |
| TtV<sub>o</sub>V<sub>1</sub> | 5.4 ± 1.3 | 0.3 ± 0.09 | 35 – 45 | 290 | 25 | (16) |

### Kinetic contributions of ΔpNa and Δψ to ATP synthesis

We next investigated contributions of Δψ and ΔpNa to the ATP synthesis rate of EhV_o_V_1_ by changing the ΔpNa or Δψ under constant Δψ (77.0 or 78.0 mV) or 2.3(*k*_B_*T*/*e*)·ΔpNa (76.5 or 76.9 mV), respectively (Figure 3A, left and right, respectively and Figure S2). The experiment was designed to compare the impacts of Δψ and ΔpNa under nearly equivalent total *smf* values ranging from ∼77 to ∼150 mV. In this series of experiments, [ADP] and [Pi] were set at 0.5 mM and 25 mM, respectively. Again, to facilitate ATP synthesis, we did not add ATP intentionally but a small amount of ATP was contaminated in ADP (<0.003%) (Table S2)

**Figure 3.**
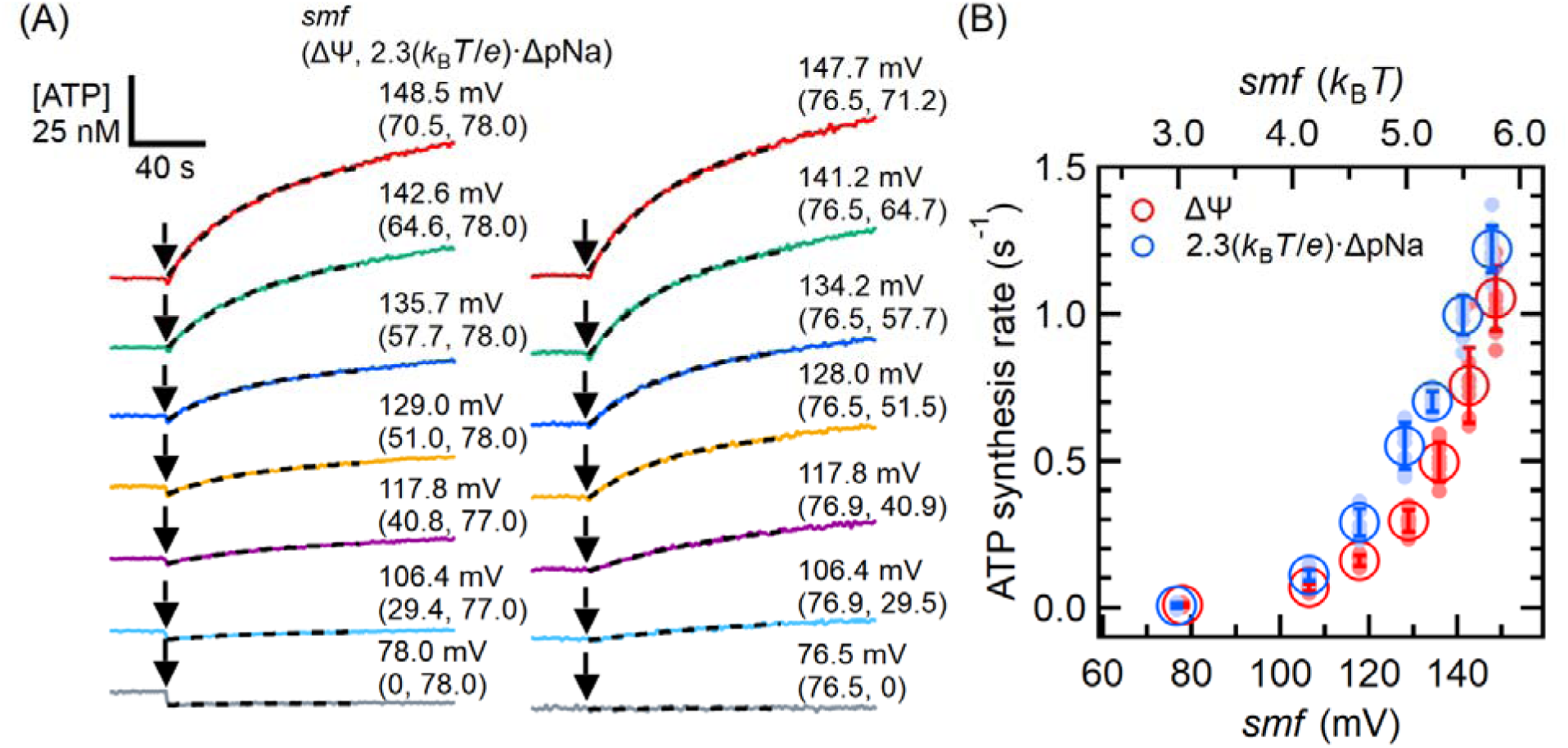
Contribution of Δψ and ΔpNa to ATP synthesis of EhV_o_V_1_. (A) Enlarged view of typical time courses of ATP synthesis at different *smf*. The entire time courses are shown in Figure S2. The reaction was initiated by adding PL (black arrows). [Pi] and [ADP] were 100 and 0.5 mM, respectively. ATP was not added but contaminated in ADP (<0.003%) (Table S2). In the left panel, Δψ was varied from 0 to 70.5 mV under 2.3(*k*_B_*T*/*e*)·ΔpNa of 77.0 or 78.0 mV. In the right panel, 2.3(*k*_B_*T*/*e*)·ΔpNa was varied from 0 to 71.2 mV under Δψ of 76.5 or 76.9 mV. The black dashed lines represent the fitting with a single exponential function for Δt = 100 sec after the addition of PL. (B) Δψ (red) and 2.3(*k*_B_*T*/*e*)·ΔpNa (blue) dependences of ATP synthesis rate. Closed and open circles represent the individual measurements (N ≥ 8) and mean values, respectively. Error bars indicate standard deviations.

As results, ATP synthesis was not detected at the lowest *smf* values, 2.3(*k_B_T*/*e*)·ΔpNa of 78.0 mV or Δψ of 76.5 mV alone (Figure 3A, bottom). However, once the total *smf* exceeded these values, ATP synthesis was observed, indicating that the threshold *smf* for ATP synthesis lies in this range. As the *smf* increased, ATP synthesis rate increased, regardless of whether Δψ or ΔpNa increased. Figure 3B shows the Δψ (red) and ΔpNa (blue) dependences of the ATP synthesis rate. Although ΔpNa exhibited a slightly larger impact than Δψ, ATP synthesis rate increased as Δψ or ΔpNa increased, suggesting nearly equivalent kinetic contributions of Δψ and ΔpNa to the ATP synthesis of EhV_o_V_1_.

#### Thermodynamic efficiency at the equilibrium points between ATP synthesis and hydrolysis

We then investigated the equilibrium points where ATP synthesis and hydrolysis are balanced. According to Equation 1 and 2, the direction of ATP synthesis/hydrolysis reaction is determined by the magnitudes of *K*_(ATP)_ and *smf*. To determine the equilibrium points under various *K*_(ATP)_ and *smf* conditions, we conducted experiments with ATP at 25 nM, Pi at 9.95 mM, and ADP varying from 20 to 80 µM, under *smf* conditions ranging from 93.5 to 132.6 mV (Table S3). Figure 4A-D and Figure S3 show the time courses of the luminescence intensity or [ATP] under these conditions. We observed ATP synthesis at high *smf* and ATP hydrolysis at low *smf*, as indicated by positive and negative slopes, respectively.

**Figure 4.**
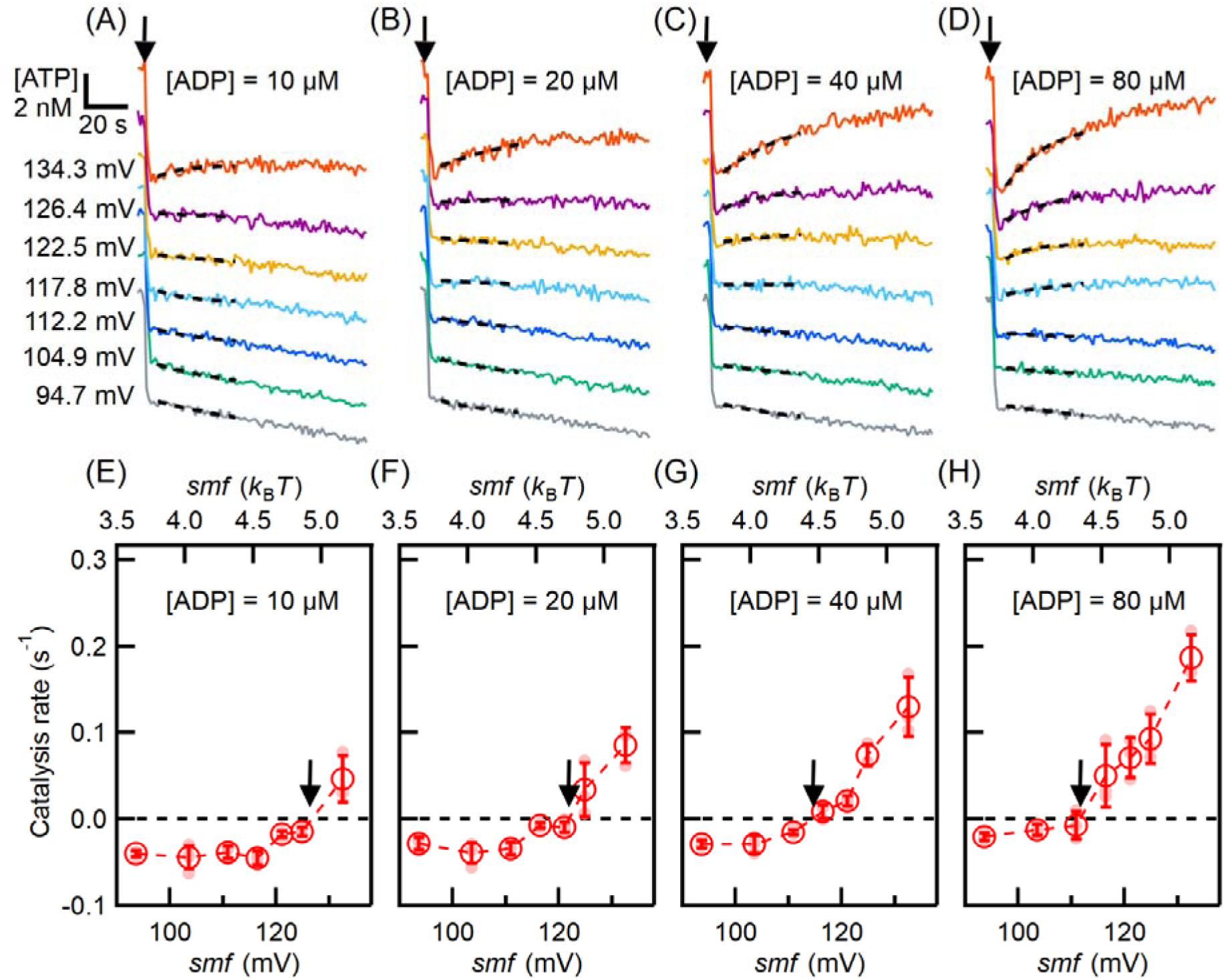
Determination of the equilibrium points between ATP synthesis and hydrolysis. (A-D) Typical time courses of ATP synthesis and hydrolysis at different *smf* (93.5 - 132.6 mV). Reaction was initiated by adding PLs as indicated by black arrow. The black dashed lines represent the fitting results obtained by fitting with a single exponential function for Δt = 35 sec after the addition of PLs. The entire time courses are shown in Figure S3. (E-H) *smf* dependence of ATP synthesis and hydrolysis rates. Black arrows indicate equilibrium points obtained by linear fitting between two data points across the catalysis rate of zero. The reaction solution contains 25 nM ATP, 9.95 mM Pi and ADP at 10 μM (A) and E), 20 μM (B and F), 40 μM (C and G), and 80 μM (D and H), respectively (Table S3).

The calculated catalysis rates were plotted against *smf* (Figure 4E-H). The positive and negative values correspond to ATP synthesis and hydrolysis rates, respectively. At low *smf*, EhV_o_V_1_ showed nearly constant ATP hydrolysis rates, presumably because ATP binding is the rate-limiting at 25 nM ATP. At high *smf*, EhV_o_V_1_ showed *smf*-dependent changes in the rates from ATP hydrolysis to synthesis. Then, we determined *smf*_eq_ (*smf* at which the net rate of ATP synthesis/hydrolysis is zero, black arrows) by linearly interpolating the two data points crossing the catalysis rate of zero. At this equilibrium point, Δ*G*’ in Equation 1 is zero and Equation 1 can be transformed into following equation:

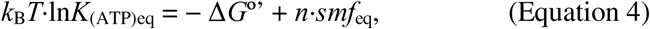

where *K*_(ATP)eq_ is the chemical potential of ATP at the equilibrium point. Based on Equation 4, *k*_B_*T*·ln*K*_(ATP)eq_ was plotted as a function of *smf*_eq_ (Figure 5) and fitted with a linear function. Slope and y-intercept of the fitted line correspond to the ion/ATP ratio *n* and the standard Gibbs free energy of ATP synthesis Δ*G*°’, respectively, and *n* of 3.2 ± 0.4 (fitted value ± SE of the fit) and Δ*G*°’ of 17.0 ± 1.7 *k*_B_*T* (42.2 ± 4.2 kJ/mol) were obtained. The value of the Δ*G*°’ was similar to those (36∼39 kJ/mol) previously reported for other ATP synthases (10,37–39), validating our experimental system. The experimentally determined value of the *n* (3.2) showed excellent agreement with that (10/3 = 3.3) expected from the structural composition of EhV_o_V_1_ which has 3 ATP catalytic sites in EhV_1_ and 10 Na^+^ binding sites in the c-ring of EhV_o_ (23–28).

**Figure 5.**
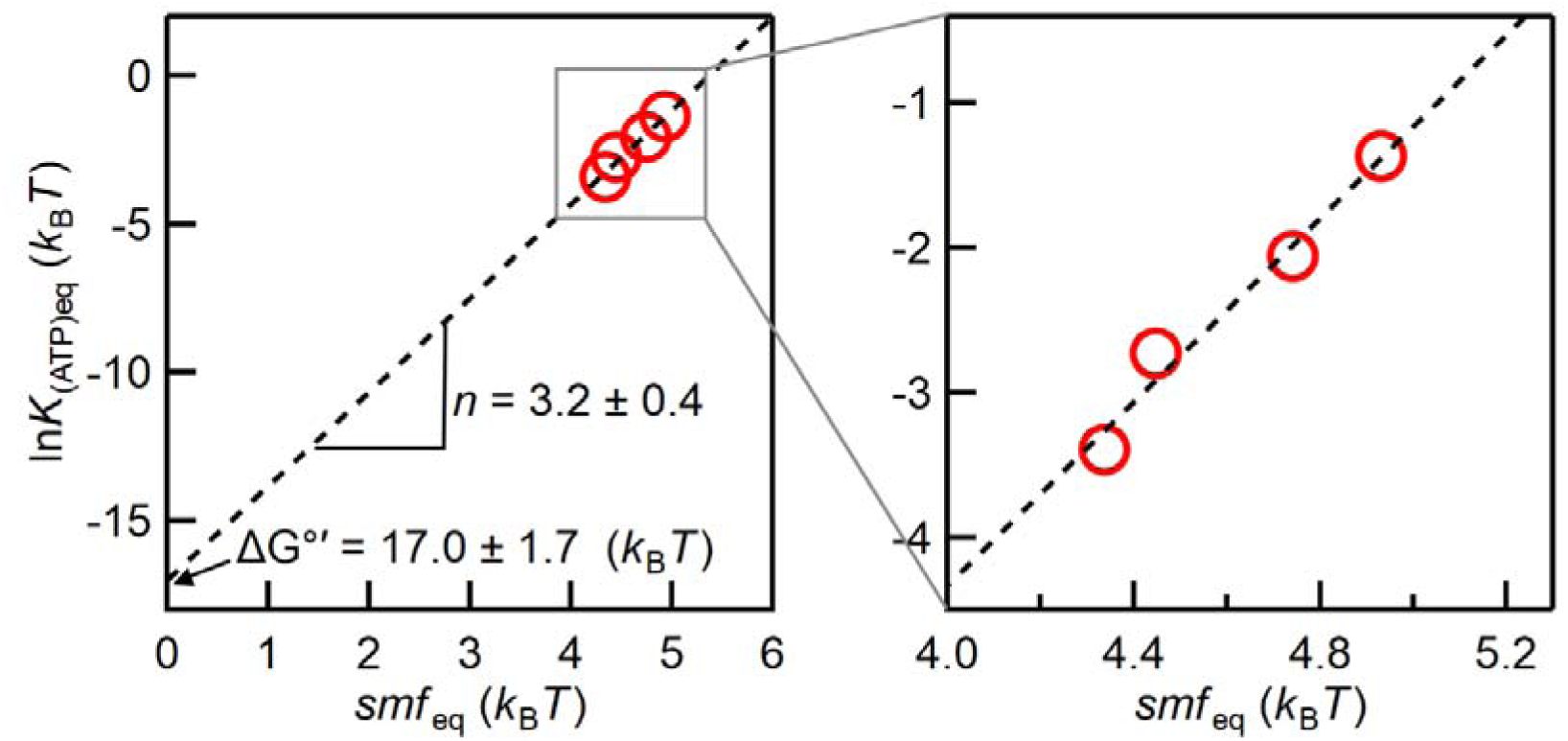
Relationship between *k*_B_*T*·ln*K*_(ATP)eq_ and *smf*_eq_ at the equilibrium points. The dashed line shows the fitting with a linear function. The slope and extrapolated y-intercept indicate the Na^+^/ATP ratio *n* (3.2 ± 0.4, fitted value ± S.E. of fitting) and the standard Gibbs free energy of ATP synthesis Δ*G*°’ (17.0 ± 1.7 *k*_B_*T* or 42.0 ± 4.2 kJ/mol), respectively.

## Discussion

The present study demonstrated that EhV_o_V_1_ is capable of synthesizing ATP driven by *smf*, despite its physiological function as an ATP-driven Na^+^ pump. The obtained values of the *K*_m_ for ADP (21 µM) and Pi (2.1 mM) during ATP synthesis of EhV_o_V_1_ are comparable to those of other ATP synthases (Table 1). The comparable values of *K*_m_ indicate similar substrate affinities during ATP synthesis across different rotary ATPases, irrespective of their primary physiological functions. The conserved properties of kinetic parameters may reflect a common evolutionary origin of these rotary ATPases (40, 41).

In particular, relatively low *K*_m_ value for Pi (high affinity of Pi) during ATP synthesis of EhV_o_V_1_ comparable to other ATP synthases is noteworthy (Table 1). Our previous single-molecule study on isolated EhV_1_ revealed low affinity of Pi and chemo-mechanical coupling scheme during rotation driven by ATP hydrolysis (30). In this scheme, Pi is released immediately after the hydrolysis of ATP into ADP and Pi without rotation, followed by release of ADP after 40° rotation. Low affinity of Pi for EhV_1_ is also supported by structural studies and MD simulation (24, 42). In contrast, F_1_ isolated from thermophilic *Bacillus* PS3 F_o_F_1_ (TF_1_), which physiologically functions as an ATP synthase, exhibits opposite order of product release during ATP-driven rotation in which ADP is released first and Pi is released second at the different rotation angles (43). This order in TF_1_ has been proposed to be favorable for ATP synthesis under physiological high [ATP] and low [ADP] environment, because Pi bound to the catalytic site would prevent ATP binding and facilitate ADP binding. Our current results with EhV_o_V_1_ suggest that the order of ADP and Pi release (or the order of ADP and Pi binding in ATP synthesis reaction) itself is not critical for the ATP synthesis capability of rotary ATPases. Our results also suggest dynamic modulation of the substrate (product) affinity depending on the rotation angle and direction, as demonstrated with TF_1_ (44). Further studies of various rotary ATPases would reveal how they have acquired functional diversity during their evolution.

In EhV_o_V_1_, both ΔpNa and Δψ contribute to driving ATP synthesis, with ΔpNa having a slightly larger contribution (Figure 3). This property is similar to Na^+^-transporting A-ATPases from *Eubacterium callanderi* and *Acerobacterium woodii* and H^+^-transporting TF_o_F_1_ (45, 46). In contrast, reports on H^+^-transporting *E. coli* F_o_F_1_ and Na^+^-transporting *Propionigenium modestum* F_o_F_1_ suggest that only one component effectively acts as the driving force (14, 45). The difference in kinetic contribution appears to be a specific feature of each rotary ATPase, independent of transporting ion species or physiological functions. Further structural studies during ATP synthesis under precisely controlled ΔpNa and Δψ are necessary to understand why these rotary ATPases show different ΔpNa and Δψ dependences. Meanwhile, because other factors such as lipid composition of the liposome and buffer and salt compositions of the assay solution may affect their ATP synthesis activities (45), the results of *in vitro* experiment should be carefully interpreted considering their physiological environments.

The nearly identical Na^+^/ATP ratios between experimental value (3.2, Figure 5) and structural prediction (3.3) provides an evidence for tight coupling between ATP synthesis/hydrolysis and Na^+^ transport in EhV_o_V_1_, consistent with our previous single-molecule study of ATP-driven rotation of EhV_o_V_1_ rate-limited by Na^+^ transport (31). In other words, EhV_o_V_1_ operates with high thermodynamic efficiency at the equilibrium point, with minimal energy loss or uncoupling. Similar excellent agreements in ion/ATP ratios have been reported for TtV_o_V_1_ and TF_o_F_1_ (10, 37), suggesting that tight coupling between ATP synthesis/hydrolysis and ion transport is a fundamental feature of rotary ATPases regardless of their physiological functions.

On the other hand, several studies have reported significant differences in the *n* values between biochemical experiments and structural predictions for ATP synthases from chloroplast, *E. coli*, and yeast (38, 39). This apparent discrepancy may reflect the different mechanisms among different rotary ATPases, or may stem from the difficulties of accurately controlling the *imf*. Factors such as unintended ion contaminations and/or ion leakages across the lipid membrane would affect the results. The use of Na^+^, which exhibits significantly lower membrane permeability than H^+^ (47), has an advantage to control the *imf* precisely.

Despite the ATP synthesis capability of EhV_o_V_1_ demonstrated *in vitro*, the physiological function appears to be limited to ATP-driven Na^+^ pumping *in vivo*. In the intestines where [Na^+^] is ∼100 mM (48, 49), *E. hirae* usually maintains Na^+^ homeostasis (∼10 mM intracellular concentration) using H^+^/Na^+^ antiporters driven by the proton motive force (*pmf*) (50, 51). When pH in the intestine increases and *pmf* generation becomes insufficient, EhV_o_V_1_ expression is induced to pump out Na^+^ using ATP hydrolysis instead of the H^+^/Na^+^ antiporters. Considering the relatively low ΔpNa of ∼1 (log_10_[100 mM/10 mM]) and high intracellular [ATP] at millimolar level, ATP synthesis of EhV_o_V_1_ would be thermodynamically unfavorable (52). Furthermore, interestingly, EhV_o_V_1_ does not have any known regulatory mechanisms that inhibit ATP hydrolysis such as ADP-Mg^2+^ inhibition (53, 54), subunit-specific inhibition (55–59), inhibition by endogenous regulatory peptide (60–62), and reversible disassembly of V_1_ and V_o_ (6, 63, 64), which are commonly found in many ATP synthases and V-ATPases. Therefore, EhV_o_V_1_ is likely specialized as ATP-driven Na^+^ pump rather than ATP synthase *in vivo*.

## Experimental procedures

### Expression and purification of EhV_o_V_1_

The expression and purification of EhV_o_V_1_ were performed according to a previously described method (31, 32). Briefly, inverted membranes prepared from *E. coli* (C41(DE3)) expressing EhV_o_V_1_ with histidine-tags in the c-subunits were solubilized using 2% *n*-dodecyl β-_D_-maltoside (DDM). The solubilized suspension was then applied to a nickel-nitrilotriacetic acid agarose (Ni-NTA agarose, QIAGEN) column. The EhV_o_V_1_ complex was eluted using a buffer containing 50 mM potassium phosphate (pH7.0), 5 mM MgCl_2_, 100 mM NaCl, 10% glycerol, and 0.05% DDM. After the elution, the complex was concentrated and subjected to size-exclusion chromatography using Supderdex 200 Increase 10/300 column (Cytiva). The column was equilibrated with a buffer containing 50 mM Tris-HCl (pH7.5), 5 mM MgCl_2_, 50 mM NaCl, 10% glycerol, and 0.05% DDM. The chromatography was performed using a fast protein liquid chromatography system (Äkta go GE Healthcare) at 4°C. The purified EhV_o_V_1_ was concentrated to approximately 20 μM and stored at −80°C until use.

### Reconstitution of EhV_o_V_1_ into liposome

L-α-Phosphatidylcholine from soybean (Type II-S, Sigma-Aldrich) was washed with acetone and suspended in reconstitution buffer (100 mM BisTris (pH7.0) containing 5 mM MgCl_2_ and 150 mM sucrose, and specific concentrations of NaCl and KCl described in Table S1-S3) at 40 mg/mL. The suspension underwent three freeze-thaw cycles. After ultracentrifugation (150,000×g for 90 min at 4°C), the pellet was resuspended in reconstitution buffer and subjected to another freeze-thaw cycle. This procedure was repeated three times to remove unintended Na^+^ and K^+^ from the lipids. The liposome was stored at −80°C until use. Proteo-liposome (PL) was prepared by mixing liposome with purified EhV_o_V_1_ at a volume ratio of 99:1. The mixture was gently mixed by inverting the tube 20 times and subjected to one freeze-thaw cycle. The final concentration of EhV_o_V_1_ in the PL was 0.2 μM. The prepared PL was stored at −80°C until use.

### Measurement of [Na^+^] and [K^+^] unintendedly contained in samples

[Na^+^] and [K^+^] unintendedly contained in the Type II-S lipid and a luciferin/luciferase reagent (ATP bioluminescence assay kit CLS II, Roche) were estimated using inductively coupled plasma optical emission spectrometer (ICP-OES, Agilent Technologies). The flow rates of the plasma and assist gas (argon) were 14 and 12 L/min, respectively. The emission intensities were measured at λ = 589.592 and 766.491 nm for sodium and potassium, respectively. Samples were prepared as follows: luciferin/luciferase reagent was dissolved in ultra-pure water (18 mg/mL), Type II-S lipid was suspended in ultra-pure water (40 mg/mL) and liposomes were prepared by the above method (40 mg/mL). For calibration, each sample was supplemented with a sodium or potassium standard solution (FUJIFILM Wako). The concentrations of sodium and potassium standard solutions were adjusted between 0.01 and 21.7 mM and between 0.639 and 63.9 mM, respectively, with specific ranges for Type II-S lipid and a luciferin/luciferase reagent samples. After addition of the standard solution, samples were further diluted with ultra-pure water to bring them within the calibration range, and the final concentrations were calculated considering the dilution factors. Each measurement was performed in triplicate, and the average values were used for analysis.

### Hexokinase treatment with ADP

Commercial ADP (117105, Merck) was treated with hexokinase (H4502, Sigma) to reduce ATP contamination as described in a previous study (14). Briefly, 970 µL of ADP solution (∼200 mM) was incubated with 1.3 mg hexokinase, 20 mM glucose, and 5 mM MgSO_4_ at 25°C for 30 minutes. ADP was then purified by gel filtration (NAP10, Cytiva) and stored at −80°C until use. The treated ADP was used for all ATP synthesis measurements.

### Measurement of ATP synthesis/hydrolysis activity

ATP synthesis and hydrolysis activities of EhV_o_V_1_ reconstituted into liposome were measured using a luminometer (AB-2270, ATTO) and the luciferin/luciferase reagent (ATP bioluminescence assay kit CLS II, Roche). ATP-dependent light emission from luciferin catalyzed by luciferase was monitored (10, 14, 15). The reaction mixture was prepared in a 1.5 mL tube containing 890 µL of observation buffer (100 mM BisTris (pH7.0) containing 5 mM MgCl_2_ and 150 mM sucrose, and specific concentrations of NaCl, KCl, and phosphate as detailed in Table S1-S3), 10 µL of valinomycin (final concentration: 83 nM), and 100 µL of luciferin/luciferase reagent (final concentration: 1.5 mg/mL). The tube was placed in the luminometer, and the measurement was initiated. At 10 seconds, 100 µL of hexokinase-treated ADP was added at the desired concentration, ranging from 0.001 to 0.5 mM. At 60 seconds, 100 µL of PL sample was introduced. The PL sample was prepared immediately before the measurement by mixing 5 µL of PL suspension with 95 µL of observation buffer, resulting in a 20-fold dilution. Finally, at 240 seconds, 100 µL of ATP (final concentration: 200 nM) was added for calibration. The total measurement time was 300 seconds.

For experiments determining the equilibrium point of ATP synthesis and hydrolysis reactions, the order and timing of additions were modified as follows. ADP (final concentrations: 10, 20, 40, or 80 µM) was added at 10 seconds, ATP (final concentration: 25 nM) at 60 seconds, and the diluted PL sample at 100 seconds. The total measurement time for these experiments was 200 seconds.

The rate of ATP synthesis/hydrolysis was determined by fitting the luminescence intensity change after the addition of PL with a single exponential function and calculating the initial slope of the fitted curve. All measurements were performed in more than triplicate at 25°C.

The *smf* for each experimental condition was calculated by considering both the intentionally added [Na^+^] and [K^+^] and the unintended contamination of these ions in the Type II-S lipid and a luciferin/luciferase reagent, as measured by ICP-OES. This approach ensured accurate quantification of the actual *smf*.

## Data availability

All data supporting this work are available from the corresponding authors upon reasonable request.

## Supporting Information

This article contains supporting information.

## Acknowledgements

We would like to thank Dr. Takeshi Murata and Dr. Takanori Harashima for their helpful discussions. A part of this work was performed with the aid of Instrument Center, Institute for Molecular Science.

## Author contributions

**Akihiro Otomo:** Conceptualization, Methodology, Validation, Formal analysis, Investigation, Resources, Data Curation, Writing - Original Draft, Visualization, Supervision, Project administration, Funding acquisition. **Lucy Gao Hui Zhu:** Validation, Investigation. **Yasuko Okuni:** Resources. **Mayuko Yamamoto:** Validation, Investigation, Resources. **Ryota Iino:** Conceptualization, Methodology, Validation, Writing - Review & Editing, Visualization, Supervision, Project administration, Funding acquisition.

## Funding

This work was supported by JSPS KAKENHI, Japan Society for the Promotion of Science Grants-in-Aid for Scientific Research (JP18H0524, JP21H02454, and JP24K01996 to R.I., JP21K15060 and JP24K17026 to A.O.) and in part by Tatematsu Foundation to A.O.

## Conflict of Interest

The authors declare that they have no known competing financial interests or personal relationships that could have appeared to influence the work reported in this paper.

## Abbreviations

The abbreviations used are: ATP, adenosine triphosphate; ADP, adenosine diphosphate; Pi, inorganic phosphate; PL, proteoliposome; imf, ion motive force; smf, sodium motive force; pmf, proton motive force; *E. hirae*, *Enterococcus hirae*; EhV_o_V_1_, V_o_V_1_ from *Enterococcus hirae*; TF_o_F_1_, F_o_F_1_ from thermophilic *Bacillus* PS3

## Notes

### Competing Interest Statement

The authors have declared no competing interest.

