## Supporting Information for "ATP synthesis of *Enterococcus hirae* V-ATPase driven by sodium motive force"

**Table S1**. Concentrations of key components for the measurements in Figure 2

| | [ADP] dependence | [Pi] dependence |
|---|---|---|
| $[Na^+]_{in}$ (mM) | 200 | 200 |
| $[Na^+]_{out}$ (mM) | 2.3 | 2.3 |
| ΔpNa (mV) | 114.6 | 114.6 |
| $[K^+]_{in}$ (mM) | 1.1 | 1.1 |
| $[K^+]_{out}$ (mM) | 454.4 | 454.4 |
| Δψ (mV) | 154.7 | 154.7 |
| ATP (nM) | 0* | 0* |
| ADP (mM) | 0.001 - 0.5 | 0.5 |
| Pi (mM) | 74 | 0.2 - 100 |

*ATP was not intentionally added but contaminated in ADP (<0.003%).

**Table S2**. Concentrations of key components for the measurements in Figure 3 and Figure S2

| | Δψ dependence | | | | | | |
|---|---|---|---|---|---|---|---|
| $[Na^+]_{in}$ (mM) | 250 | 250 | 250 | 250 | 250 | 250 | 250 |
| $[Na^+]_{out}$ (mM) | 12.0 | 12.5 | 12.5 | 12.0 | 12.0 | 12.0 | 12.0 |
| ΔpNa (mV) | 78.0 | 77.0 | 77.0 | 78.0 | 78.0 | 78.0 | 78.0 |
| $[K^+]_{in}$ (mM) | 6.2 | 12.9 | 12.9 | 6.2 | 6.2 | 6.2 | 6.2 |
| $[K^+]_{out}$ (mM) | 45.1 | 40.6 | 63.2 | 45.1 | 58.6 | 76.7 | 96.6 |
| Δψ (mV) | 0* | 29.4 | 40.8 | 51.0 | 57.7 | 64.6 | 70.5 |
| ATP (nM) | 0** | | | | | | |
| ADP (mM) | 0.5 | | | | | | |
| Pi (mM) | 100 | | | | | | |

| | ΔpNa dependence | | | | | | |
|---|---|---|---|---|---|---|---|
| $[Na^+]_{in}$ (mM) | 250 | 250 | 250 | 250 | 250 | 250 | 250 |
| $[Na^+]_{out}$ (mM) | 250.7 | 78.9 | 50.9 | 33.7 | 26.5 | 20.2 | 15.6 |
| ΔpNa (mV) | 0 | 29.5 | 40.9 | 51.5 | 57.7 | 64.7 | 71.2 |
| $[K^+]_{in}$ (mM) | 6.2 | 12.9 | 12.9 | 6.2 | 6.2 | 6.2 | 6.2 |
| $[K^+]_{in}$ (mM) | 122 | 257.6 | 257.6 | 122 | 122 | 122 | 122 |
| Δψ (mV) | 76.5 | 76.9 | 76.9 | 76.5 | 76.5 | 76.5 | 76.5 |
| ATP (nM) | 0** | | | | | | |
| ADP (mM) | 0.5 | | | | | | |
| Pi (mM) | 100 | | | | | | |

*Valinomycin was not added so Δψ is practically zero.

**ATP was not intentionally added but contaminated in ADP (<0.003%).

**Table S3**. Concentrations of key components for the measurements in Figure 4 and Figure S3

| | | | | | | | |
|---|---|---|---|---|---|---|---|
| $[Na^+]_{in}$ (mM) | 250 | 250 | 250 | 250 | 250 | 250 | 250 |
| $[Na^+]_{out}$ (mM) | 20.2 | 20.2 | 20.2 | 20.2 | 20.2 | 20.2 | 20.2 |
| ΔpNa (mV) | 64.6 | 64.6 | 64.6 | 64.6 | 64.6 | 64.6 | 64.6 |
| $[K^+]_{in}$ (mM) | 6.2 | 6.2 | 6.2 | 6.2 | 6.2 | 6.2 | 6.2 |
| $[K^+]_{out}$ (mM) | 19.1 | 28.2 | 37.4 | 46.5 | 55.6 | 64.7 | 87.5 |
| Δψ (mV) | 28.9 | 38.9 | 46.2 | 51.8 | 56.4 | 60.2 | 68.0 |
| ATP (nM) | 25 | | | | | | |
| ADP (mM) | 0.01 - 0.08 | | | | | | |
| Pi (mM) | 9.95 | | | | | | |

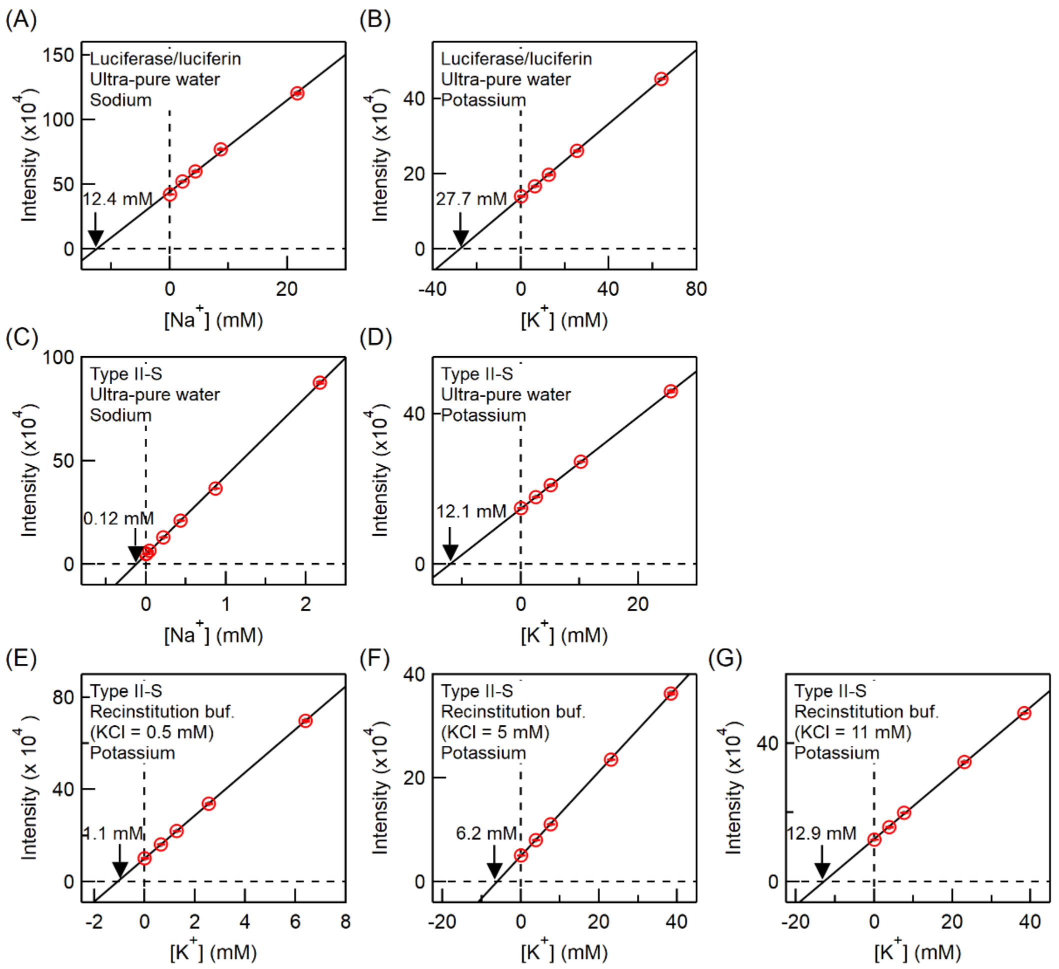


**Figure S1**. Quantitative analysis of $Na^+$ and $K^+$ in luciferin/luciferase reagent and Type II-S lipid by ICP-OES using standard addition method. The X-intercepts (indicated by black arrows) obtained by extrapolation of the calibration curves give the concentration of analyte in the samples. (A and B) $Na^+$ and $K^+$ concentrations in 18 mg/mL of the luciferin/luciferase reagent dissolved in ultra-pure water. (C and D) $Na^+$ and $K^+$ concentrations in 40 mg/mL of the Type II-S lipid suspended in ultra-pure water. (E to G) $K^+$ concentrations in 40 mg/mL of Type II-S lipids suspended in reconstitution buffers prepared with 0.5, 5 mM, and 11 mM of KCl, respectively.

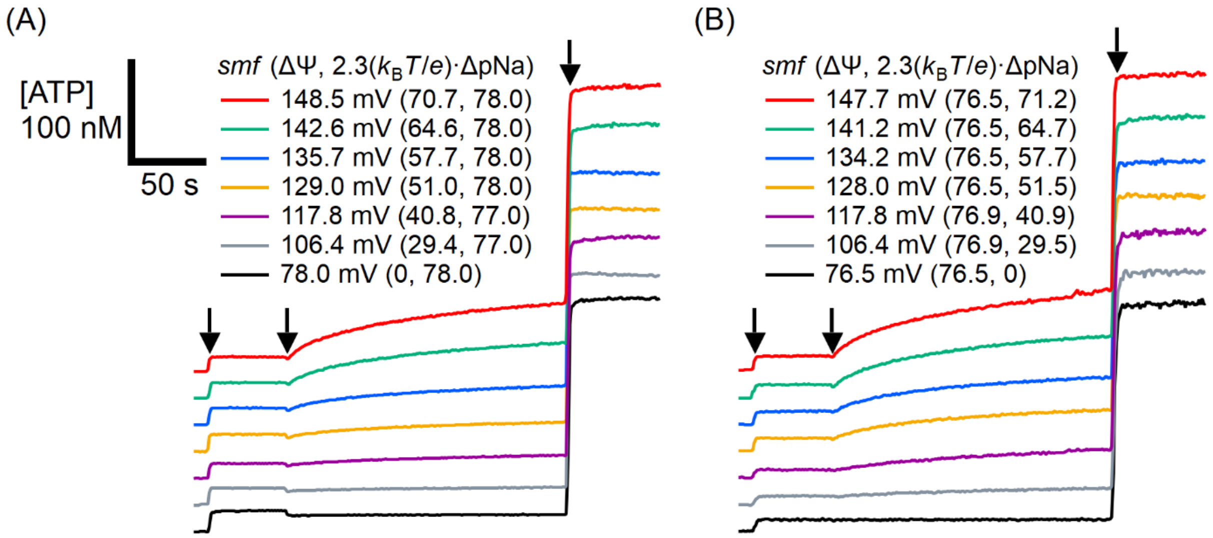


**Figure S2**. Typical time courses of ATP synthesis at different *smf*. ADP (final concentration: 0.5 mM), PL, and ATP (final concentration: 200 nM) were added at 10, 60, and 240 sec, respectively, as indicated by black arrows. (A) Δψ dependence under $2.3(k_B T/e)\cdot\Delta pNa$ of 77.0 or 78.0 mV. (B) ΔpNa dependence under Δψ of 76.5 or 76.9 mV.

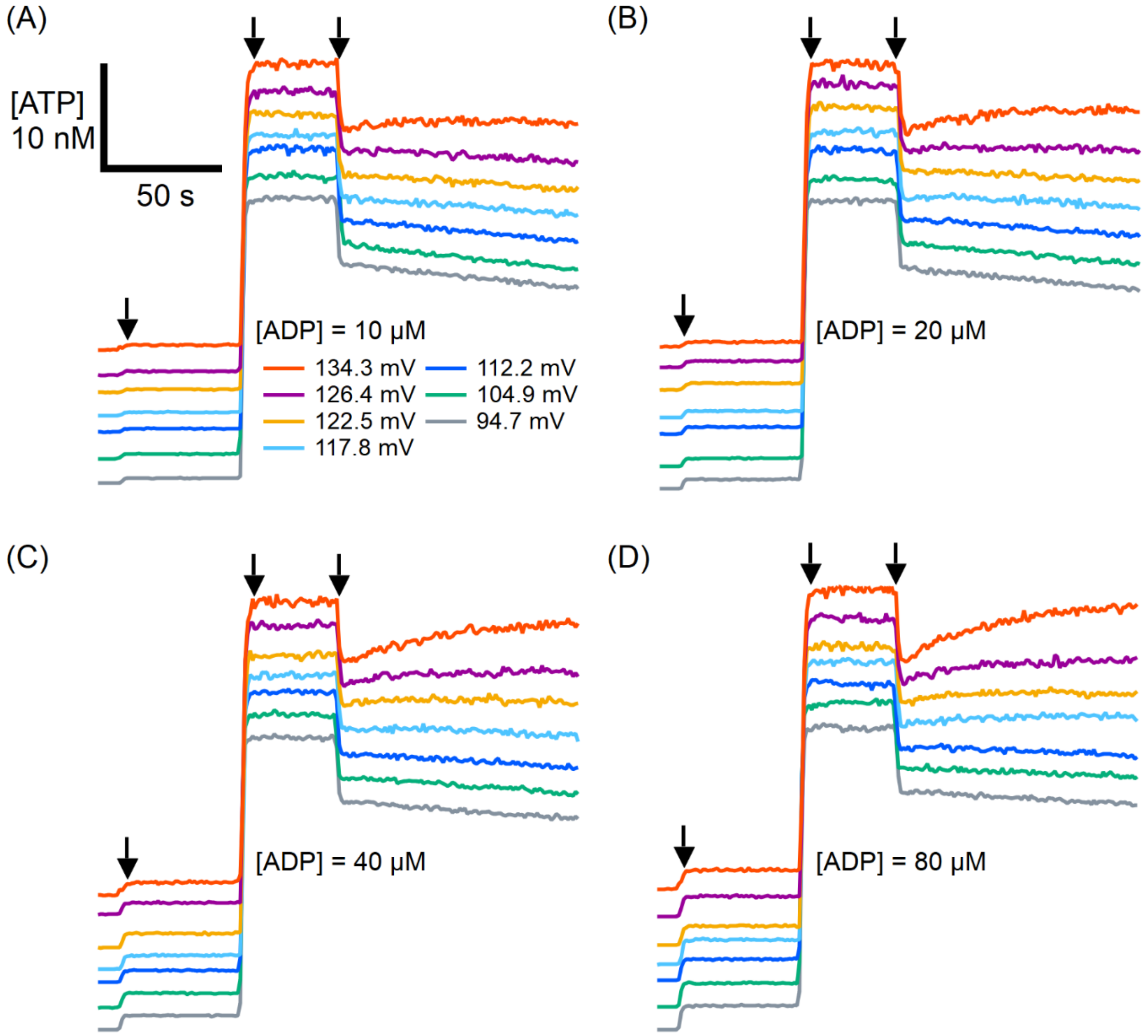


**Figure S3**. Typical time courses of ATP synthesis and hydrolysis at different *smf* (94.7 – 134.3 mV). ADP (10 - 80 μM), ATP (25 nM), and PL were added at 10, 60, and 100 sec, respectively, as indicated by black arrows. ADP concentrations were 10 μM (A), 20 μM (B), 40 μM (C), and 80 μM (D). Pi concentration was 9.95 mM for all conditions.
